# Scalable Nonlinear Programming Framework for Parameter Estimation in Dynamic Biological System Models

**DOI:** 10.1101/410688

**Authors:** Sungho Shin, Ophelia Venturelli, Victor M. Zavala

## Abstract

We present a nonlinear programming (NLP) framework for the scalable solution of parameter estimation problems that arise in dynamic modeling of biological systems. Such problems are computationally challenging because they often involve highly nonlinear and stiff differential equations as well as many experimental data sets and parameters. The proposed framework uses cutting-edge modeling and solution tools which are computationally efficient, robust, and easy-to-use. Specifically, our framework uses a time discretization approach that: i) avoids repetitive simulations of the dynamic model, ii) enables fully algebraic model implementations and computation of derivatives, and iii) enables the use of computationally efficient nonlinear interior point solvers that exploit sparse and structured linear algebra techniques. We demonstrate these capabilities by solving estimation problems for synthetic human gut microbiome community models. We show that an instance with 156 parameters, 144 differential equations, and 1,704 experimental data points can be solved in less than 3 minutes using our proposed framework (while an off-the-shelf simulation-based solution framework requires over 7 hours). We also create large instances to show that the proposed framework is scalable and can solve problems with up to 2,352 parameters, 2,304 differential equations, and 20,352 data points in less than 15 minutes. Competing methods reported in the computational biology literature cannot address problems of this level of complexity. The proposed framework is flexible, can be broadly applied to dynamic models of biological systems, and enables the implementation of sophisticated estimation techniques to quantify parameter uncertainty, to diagnose observability/uniqueness issues, to perform model selection, and to handle outliers.

**Author summary:** Constructing and validating dynamic models of biological systems spanning biomolecular networks to ecological systems is a challenging problem. Here we present a scalable computational framework to rapidly infer parameters in complex dynamic models of biological systems from large-scale experimental data. The framework was applied to infer parameters of a synthetic microbial community model from large-scale time series data. We also demonstrate that this framework can be used to analyze parameter uncertainty, to diagnose whether the experimental data are sufficient to uniquely determine the parameters, to determine the model that best describes the data, and to infer parameters in the face of data outliers.

## Introduction

Dynamic modeling is essential for understanding the behavior of biological systems. Systems of interest in this domain include microbial communities and microbiome, gene regulatory networks, and metabolic pathways [1-3]. An important task that arises in modeling studies is validation against experimental data by using parameter estimation techniques. This task is computationally challenging because of the need to solve optimization problems constrained by differential equations. Challenges arise from the dimensionality, nonlinearity, and stiffness of the dynamic model, from the incomplete observation of the system states, from the need to estimate many parameters, and from the need to handle a large number of experimental data sets.

**Fig 1.**
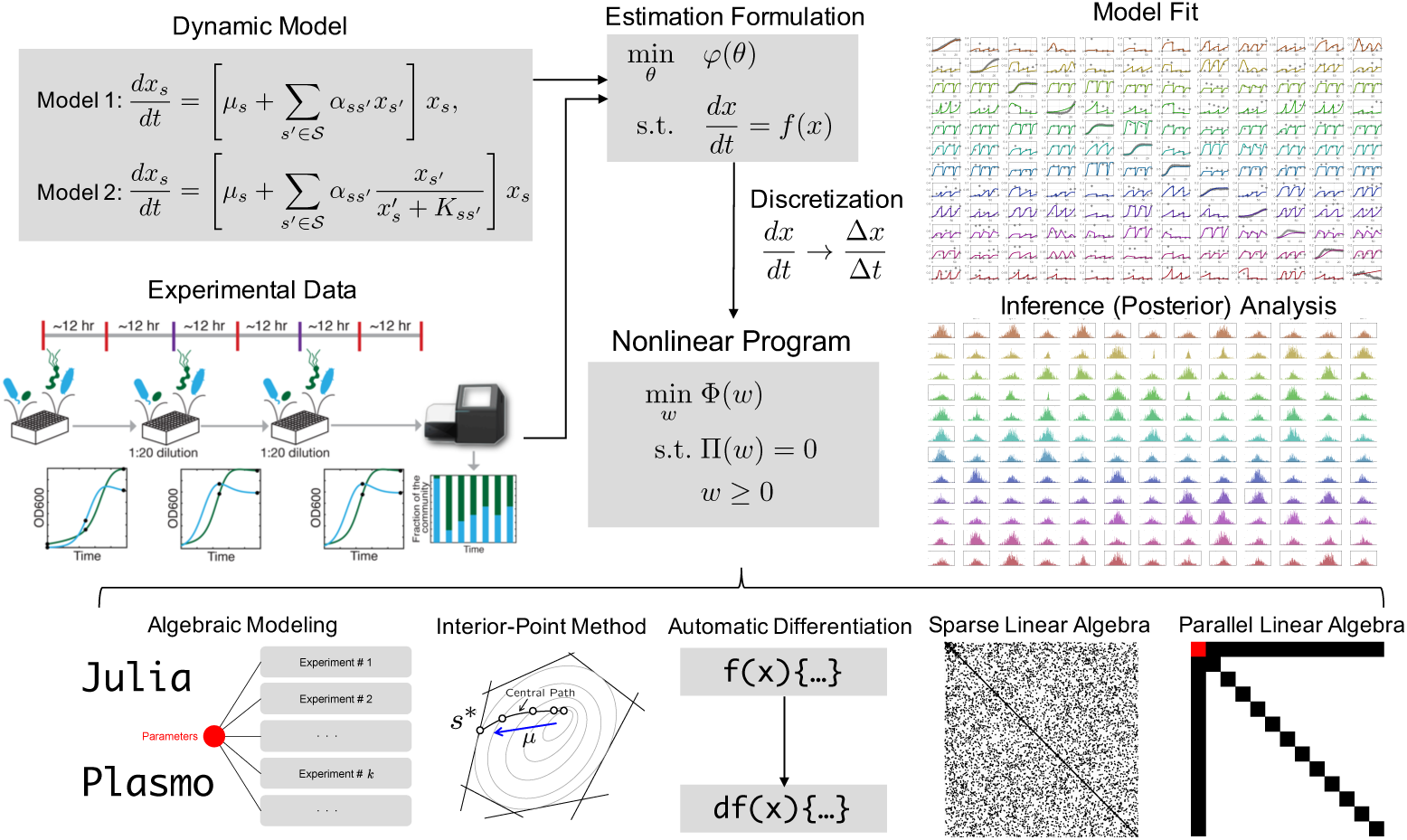
Illustration of the proposed estimation framework. Mathematical models for biological systems are often expressed as systems of differential equations with parameters that need be estimated from experimental data. We formulate the estimation problem using a maximum a posteriori (MAP) formulation. This yields optimization problems constrained by differential equations that are transformed into fully algebraic nonlinear programs by using discretization schemes. The resulting NLPs can be easily implemented in algebraic modeling languages such as `JuMP` and `Plasmo.jl` that compute derivatives automatically and that are interfaced to powerful interior-point optimization solvers that exploit sparsity and structure to achieve high computational efficiency. The proposed framework is scalable, robust, easy-to-use, and flexible. These capabilities facilitate high-level tasks such as identification of parameter observability/uniqueness issues, model selection, and uncertainty quantification.

Extensive research on solution methods for estimation problems with differential equations has been reported in the computational biology literature (see [4,5] for comprehensive reviews). These methods target maximum likelihood estimation formulations (which are derived from Bayesian principles). In these formulations, one aims to find parameters that maximize the likelihood function. The most used strategy to handle such formulations is the so-called *simulation-based* approach. Here, the idea is to perform repetitive simulations of the dynamic model at different trial parameter values to identify a set of parameters that maximizes the likelihood. The trial parameter values are updated using derivative-based or derivative-free search schemes [6-8]. While the simulation-based approach is intuitive, repetitive solutions of large dynamic models is computationally expensive and differential equation solvers can fail at trial parameter values that are non-physical or that trigger unstable responses. In addition, techniques to compute first and second order derivatives for derivative-based schemes (e.g., finite differences, forward and adjoint sensitivities) involve intrusive procedures and are often limited to first-order derivatives [7]. The need for derivatives can be bypassed by using derivative-free search schemes [9,10], which are widely popular in computational biology. Such methods include simulated annealing [11], genetic algorithms [12,13], particle swarms [14], approximate Bayesian computation [15,16], and various other methods [17,18]. Derivative-free schemes do not scale well in the number of parameters (a larger number of trial parameter values often need to be explored compared to derivative-based schemes). Moreover, second order derivative information is needed to determine if the parameter estimates are unique/observable given available experimental data [19,20]. The uniqueness/observability test of the parameter estimates is based on curvature information of the likelihood function at the solution.

Simulation-based estimation frameworks previously reported in the computational biology literature have focused on problems that usually contain less than 100 parameters [10,15,21]. To the best of our knowledge, the largest estimation problem solved using a simulation-based framework contains 3,780 data points and 1,801 parameters [7]. Such a problem was solved (to partial optimality) using a derivative-based search scheme that uses first-order derivative information (using an adjoint method) and required over 5 hours of computing time. The scalability limitations of simulation-based approaches present an important obstacle in considering models of higher fidelity, in exploiting high-throughput experimental data, in analyzing parameter uncertainties, and in implementing sophisticated techniques such as ensemble modeling.

In this work, we propose a nonlinear programming (NLP) framework for solving estimation problems with embedded dynamic models [22,23]. The framework is based on a *direct transcription* approach wherein the dynamic model is converted into a large set of algebraic equations by applying time-discretization techniques. The algebraic equations are then embedded directly as constraints in the optimization problem (a nonlinear program-NLP). The NLPs arising from time discretization are of high dimension (easily reaching hundreds of thousands to millions of variables and constraints) but are also *sparse* and *structured*. Moreover, by transforming the dynamic model into algebraic equations, it becomes possible to use automatic differentiation techniques available in modern modeling languages to compute first and second derivatives. Exploitation of sparsity and structure, together with the availability of derivative information, enable the solution of estimation problems with complex dynamic models and efficient handling of many parameters and experimental data sets.

Discretization-based estimation approaches have been widely studied in diverse fields such as chemical engineering [24-26] and aerospace engineering [27-29] (see [30] for a comprehensive review) but less so in computational biology. A major factor that has hindered wider adoption is the lack of easy-to-use computational frameworks that facilitate access to non-expert users. In this work, we demonstrate that modern modeling and solution tools can be combined to create scalable, robust, easy-to-use, and flexible frameworks. We demonstrate the benefits by solving challenging estimation problems arising in nonlinear microbial community models.

The proposed framework enables the implementation of higher level tasks such as observability analysis and uncertainty quantification. Uncertainty quantification (UQ) seeks to characterize parameter posterior distribution, which is necessary to obtain confidence levels/regions and parameter correlation information. Conventionally, UQ is performed by using second order derivative (Hessian) information of the likelihood function to construct an approximate parameter posterior covariance matrix [23,31] or by using a Markov-Chain-Monte-Carlo (MCMC) techniques [32,33]. The Hessian-based approach is scalable but it requires intrusive computation (cannot be automatically computed by the solver) and does not capture well the effect of nonlinearities and physical constraints [31]. In MCMC, one samples parameters from the prior parameter density and compares the associated model outputs with experimental data to decide whether to accept that sample or not. By repeating these accept/reject steps one can construct an approximate parameter posterior. MCMC is rather easy to implement (it is not intrusive) but, being simulation-based, also suffers from potential failures of the differential equation solver at non-physical parameter samples, it does not scale well with the number of parameters, and convergence issues might be encountered.

In this work, we propose to overcome some of these challenges by using a randomized maximum a posteriori (rMAP) framework [34-37]. This method computes approximate samples from the parameter posterior distribution by performing random perturbations on the experimental data and by re-solving the estimation problem. This allows exploration of the parameter space more efficiently compared to the MCMC scheme because each sample can be computed in parallel (MCMC is sequential). Moreover, the rMAP approach is non-intrusive, can capture nonlinear and physical constraints effects, and avoids potential failures of differential equation solvers. The proposed estimation framework is flexible and can easily accommodate advanced estimation formulations. To demonstrate this, we implement formulations that use different prior regularization schemes and *k*-max norms (the mean of a specified fraction of largest values) to mitigate large outliers [38,39].

The paper is organized as follows. The methods section provides a general form for the estimation problem under study and discusses how this can be cast as a sparse NLP by using time-discretization techniques. Furthermore, we introduce basic concepts behind NLP solvers that exploit sparsity and structures at the linear algebra level. In addition, we discuss rMAP and outlier mitigation schemes. In the results section, we demonstrate that the proposed framework can handle challenging estimation problems arising in microbial community models.

## Methods

### Estimation for Dynamic Models

We consider estimation problems of the following form:

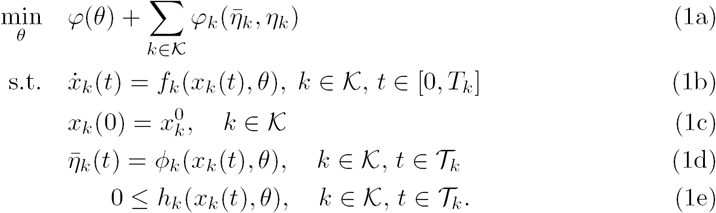

Here, *𝒦* := {0, 1… *K*} is the *set of experiments* and 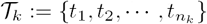 is the set of measurement (sampling) times in experiment *k ∈ 𝒦*. Time is denoted as *t ∈* [0, *T*_*k*_], where *T*_*k*_ *∈* ℝ_+_ is the duration for experiment *k ∈ 𝒦*. The variable vector 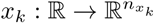 are the differential *state* time trajectories, 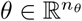 are the model *parameters*, 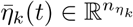 are the model predicted *outputs* with corresponding *experimental observations* 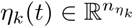 at time *t ∈ 𝒯*_*k*_ and experiment *k ∈ 𝒦*, and 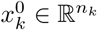 are initial conditions for experiment *k ∈ 𝒦*. For convenience in the notation, we define the output vectors 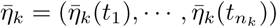 and the experimental output vectors 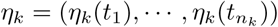 for experiment *k ∈ 𝒦* as well as the total output vector 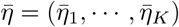 and the total experimental output vector *η* = (*η*_1_, *…, η*_*K*_). The vector function *f*_*k*_(·) denotes the dynamic model mapping, *φ*(·) and *φ*_*k*_(·) are objective function mappings, *ϕ*_*k*_(·) is the state-to-output mapping, and *h*_*k*_(·) is the constraint mapping. All the mappings are assumed to be at least twice continuously differentiable with respect to all the arguments. The estimation formulation (1a) captures all the features that we need to demonstrate the capabilities of our proposed framework. Our framework, however, can also accommodate more general features; for instance, the initial conditions (1c) can be also considered as unknown variables that need to be estimated and we can define non-additive objective functions that penalize large errors.

Problem (1) can be derived from Bayesian principles. To see this and introduce some useful notation, we start from Bayes theorem:

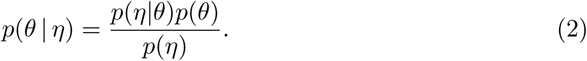

Here, *p*(*θ | η*) is the parameter posterior density (i.e., the parameter density given knowledge on the outputs), *p*(*η | θ*) is the output posterior density (i.e., the outputs given knowledge on the parameters), *p*(*θ*) is the prior density (i.e., parameter density before knowledge of the output), and *p*(*η*) is the output marginal density. In a maximum a posteriori (MAP) formulation, the goal is to find the parameters that maximize *p*(*θ | η*). Because *p*(*η*) does not depend on *θ*, this can also be achieved by maximizing *p*(*η|θ*)*p*(*θ*). Maximizing *p*(*η|θ*)*p*(*θ*) is equivalent to maximizing the log-likelihood function *L*(*θ*) = log(*p*(*θ*)) + log *p*(*η|θ*). If the outputs from the experiments *k ∈ 𝒦* are independent (which is usually the case), we have that *p*(*η|θ*) = Π_*k∈ 𝒦*_ *p*(*η*_*k*_ *| θ*) and thus:

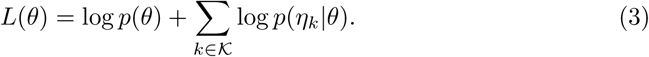

The observed outputs are random variables that are usually considered to be Gaussian and thus 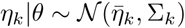, where ∑_*k*_ is the covariance matrix. The prior density *p*(*θ*) is also often assumed to be Gaussian and thus 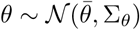, where 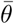 is the mean of the prior distribution and ∑_*θ*_ is its covariance. With this, we obtain:

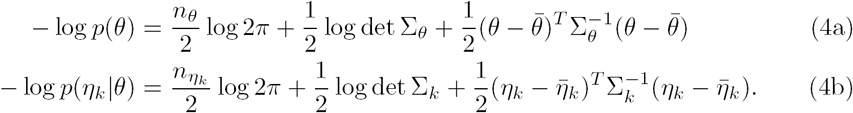

By comparing (3) with (1a) we can see that minimizing 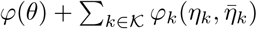 is equivalent to minimizing –*L*(*θ*). Here, the dynamic model together with the state-to-output mapping defines a *parameter-to-output mapping* of the form 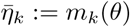. In a simulation-based estimation approach, the mapping *m*_*k*_(*θ*) is computed by simulating the dynamic model (1b)-(1c) at a trial value *θ* using a differential equation solver and by evaluating the outputs at the sampling times *t ∈ 𝒯*_*k*_ using (1d). In a discretization-based approach, the mapping *m*_*k*_(*θ*) is not computed explicitly (but we use it here as a mathematical representation that is used to explain some relevant concepts). Constraints (1e) restrict the parameter space to be explored.

After dropping all constant terms in the likelihood function we obtain:

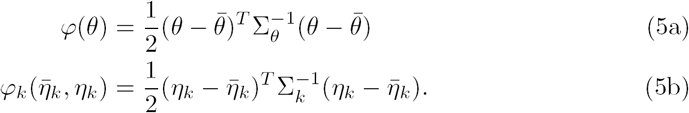

The function *ϕ*(*θ*) is usually known as the *prior* term and provides a *regularization* effect that stabilizes the solution of the estimation problem when the parameters cannot be uniquely infered from the available data [40-43]. This regularization term arises from the prior density *p*(*θ*) and provides a mechanism to encode knowledge on the parameters. Assuming that the prior density is Gaussian gives rise to a prior term that is defined by a weighted L2 norm. Recently, the machine learning community has also proposed the use of regularization terms that use L1 norms 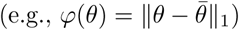. The L1 norm induces sparsity in the parameters and corresponds to assuming that the prior density is Laplacian. One can also show that an L1 norm acts as an exact penalty function and implicitly induces constraints on the parameters. Similarly, one can also use the inequality constraints *h*_*k*_(·) to directly embed physical knowledge in the MAP formulation (e.g., concentrations can only be positive).

From (5) we see that *φ*_*k*_(·) are squared error terms and thus the MAP problem minimizes the *sum* of the squared errors across all experiments *k* ∈ 𝒦. This approach offers limited control on large errors that might result from data outliers. Here, we propose to use a *k*-max norm to mitigate these issues. Our proposal is based on the observation that a *k*-max norm is equivalent to a conditional-value-at-risk (CVaR) norm [38,39,44]. The CVaR_*β*_ norm of a vector **e** = (*e*_1_, *…, e*_*K*_) with components 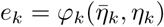 is defined as the average of the *β*-fraction of largest elements of the vector (where *β* ∈ [0, 1] is a parameter that defines the size of the fraction) [44]. One can show that, when *β* → 1, the CVaR norm is the largest fitting error and, when *β* → 0, the CVaR norm is the sum of fitting errors (as in the standard MAP formulation). A key computational property of the CVaR norm is that it can be formulated as an standard optimization problem. In particular, the MAP problem with a CVaR error norm can be expressed as:

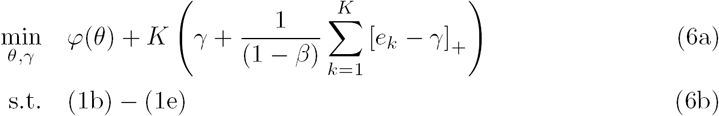

*where* [·]^+^ = max(0, *·*) is the max function and *γ* is an auxiliary variable [39].

### Nonlinear Programming Formulation

To solve the MAP problem (1) we approximate the differential equations by using a discretization scheme. This enables the use of computationally efficient NLP solvers and facilitates high-level UQ and observability monitoring tasks.

#### Time Discretization

Discretization schemes such as Euler, Runge-Kutta, and orthogonal collocation are commonly used to transform differential equations into algebraic ones. Orthogonal collocation is often preferred because accurate approximations can be obtained with few discretization points [22]. To simplify the presentation we use an implicit Euler scheme, which can be shown to be a special type of an orthogonal collocation scheme (i.e., it is a one-point Radau collocation scheme). We discretize the time domain [0, *T*_*k*_] into a set of intervals with fixed discrete-time points 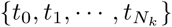 for each experiment *k ∈ 𝒦* (where *t*_0_ = 0 and 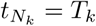). The associated index set is represented by *𝒩*_*k*_ := {1, …, *N_k_*}. By applying an implicit Euler scheme, the dynamic model (1b) is converted into a set of nonlinear algebraic equations of the form:

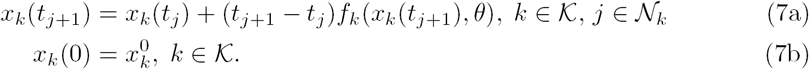

With this, we can approximate the MAP problem (1) using the NLP:

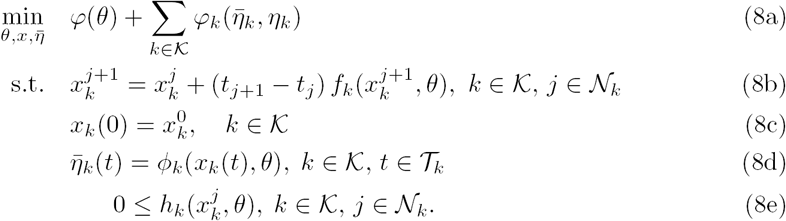

Here, we use 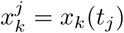 as short-hand notation to represent states at time *t*_*j*_ and experiment *k*. For convenience, we express the NLP in the following abstract form:

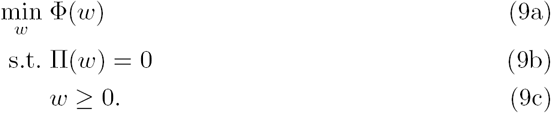

where *w ∈* ℝ^*n*^ is a large-dimensional vector containing all the discrete-time states 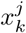, parameters *θ*, and additional auxiliary variables. The mapping Φ: ℝ^*n*^ → ℝ is the objective function and Π: ℝ^*n*^ → ℝ ^*m*^ are equality constraints that contain algebraic equations obtained fro discretization of the dynamic model and other auxiliary equations. General inequality constraints can be transformed into equality constraints and simple non-negativity bounds by using auxiliary slack variables 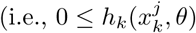 can be written as 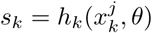 with *s*_*k*_ ≥ 0).

A useful representation of the NLP results from noticing that the parameters *θ* are the only complicating (coupling) variables across experiments *k* ∈ *𝒦.* Consequently, we can express the NLP in the *structured* from [45]:

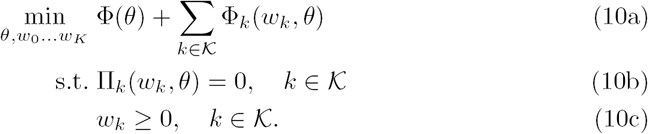

Here, the variable vector *w*_*k*_ contains all the discrete-time states and auxiliary variables of experiment *k ∈ 𝒦*, Φ(·) is the prior term, Φ_*k*_(·) is the contribution of experiment *k ∈ 𝒦* to the likelihood function, and Π_*k*_(·) contains the discretized dynamic model equations and auxiliary equations for experiment *k ∈ 𝒦*. As we discuss next, this representation can be used to derive parallel solution approaches.

#### Interior-Point Solvers

The NLPs that result from time discretization exhibit a high degree of *algebraic sparsity* (only a few variables appear in each constraint) and are highly structured. Sparsity and structure permeates down to linear algebra operations performed inside the optimization solver. This is sharp contrast to the simulation-based approach, which induces dense linear algebra operations in the space of the parameters *θ*. Most modern large-scale NLP solvers such as `Ipopt` and `Knitro` seek to exploit sparsity and structure at the linear algebra level to achieve high computational efficiency [46,47]. Interior point solvers, in particular, provide a flexible framework to do this. These solvers replace the variable bounds by using a logarithmic barrier function. In the context of NLP (9), this results in a logarithmic barrier subproblem of the form:

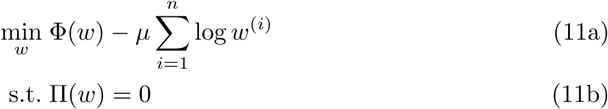

where *µ* ∈ ℝ_+_ is the so-called barrier parameter. The logarithmic term becomes large as *w*^(*i*)^ approaches the boundary of the feasible region. This ensures that variables remain in the *interior* of the feasible region (hence the origin of the term *barrier*). A key observation is that one can recover a solution of the original NLP (9) by solving a sequence of barrier problems for decreasing values of *µ* [48]. An important property of interior-point methods is that the original NLP with bounds is converted into a sequence of NLPs with equality constraints. This removes the combinatorial complexity of identifying the set of bounds that are active or inactive at the solution (a bottleneck in active-set solvers).

#### Sparse Linear Algebra

Interior-point methods enable efficient linear algebra implementations. To explain how this is done, we note that the optimality conditions of the barrier problem are given by the following set of nonlinear equations:

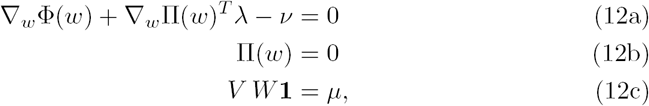

where, *λ ∈* ℝ^*m*^ are the Lagrange multipliers of the equality constraints, *v* are the Lagrange multipliers of the bound constraints, *V* = diag(*v*) and *W* = diag(*w*) are diagonal matrices, and **1** is a vector of all ones.

By applying Newton’s method to (12), we obtain the following linear algebra system:

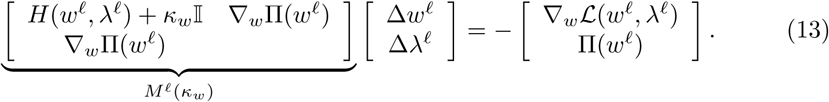

Here, *𝓁* is the Newton iteration index, Δ*w*^*𝓁*^ is the search direction for the primal variables, Δ*λ*^*𝓁*^ is the search direction for the dual variables, and 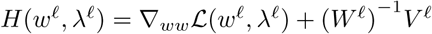 is the Hessian of the Lagrange function 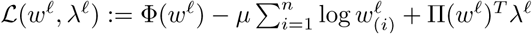. The matrix *M* ^𝓁^(*κ*_*w*_) is known as the *augmented matrix*. The Newton step computation in a simulation-based approach operates only in the space of the parameters *θ* (the states are implicitly eliminated by simulation). In the time discretization approach, the Newton search is in the space of both the discretized states and parameters (contained in the high-dimensional variable vector *w*). Interestingly, however, the augmented matrix found in typical applications is highly sparse (with less than 1% of its entries are non-zero) [23].

The constant *κ*_*w*_ *∈* ℝ_+_ is a *Hessian regularization parameter* which plays a key role in the context of parameter estimation. In particular, one can prove that the augmented matrix *M* ^𝓁^(*κ*_*w*_) is non-singular (and thus the linear algebra system has a unique solution) if and only if the reduced Hessian matrix *Z*^*T*^ *H*(*w*^𝓁^, *λ*^𝓁^)*Z* is positive definite and the Jacobian matrix *∇*_*w*_Π(*w*^𝓁^) has full row rank. Here, the matrix 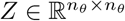 is such that its columns span the null-space of the Jacobian (i.e., ∇_*w*_Π(*w*^𝓁^)*Z* = 0). Moreover, the matrix *Z* is of the same dimension as the number of degrees of freedom (in our context this is precisely the number of parameters). When the reduced Hessian is positive definite (i.e., all its eigenvalues are positive) and the Jacobian has full row rank, one can prove that the Newton step of the primal variables Δ*w*^𝓁^ obtained from the solution of (12) is a descent direction for the objective function (i.e., (Δ*w*^𝓁^)^T^Δ_*w*_Φ(*w*^𝓁^) < 0) when the constraints are close to being satisfied (i.e., Π(*w*^𝓁^) ≈ 0). This is key because it indicates that the Newton step improves the objective function (in our context, the negative likelihood function). This property cannot be guaranteed when the reduced Hessian is not positive definite. When such a situation is encountered, one can increase the regularization parameter *κ*_*w*_ until the reduced Hessian is positive definite and a descent direction is obtained. This approach is closely connected to the Levenberg-Marquardt method used in simulation-based estimation approaches (in which one regularizes the Hessian of the negative likelihood function as – *∇* _*θθ*_*L*(*θ*^𝓁^) + *κ*_*θ*_ 𝕀) [49]. Another key observation is that, when the reduced Hessian is positive definite at the solution *w*^*\**^, the estimated parameters are *unique*. This provides an indication that the experimental data is sufficiently informative to identify the parameters uniquely (i.e., the parameters are observable). We note that using a prior term *Φ*(*θ*) in the MAP formulation has the effect of adding the positive definite matrix 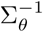 to the reduced Hessian. This artificially regularizes the problem (as is done in the Levenberg-Marquardt scheme by adding the term *κ*_*w*_𝕀). Consequently, when testing for observability/uniqueness, it is necessary to drop the prior term from the MAP formulation. Testing for observability also requires exact second order derivative information because the Hessian is needed. In the time-discretization approach, such information can be obtained directly from algebraic modeling languages. Simulation-based solution approaches often cannot check observability of the parameters (computing second derivatives using adjoint and sensitivity schemes is complicated).

Computing the eigenvalues of the reduced Hessian to check for positive definiteness is expensive. Interestingly, one can also determine if the reduced Hessian is positive definite by using inertia information of the augmented matrix *M* ^𝓁^(*κ*_*w*_). The inertia of a matrix *M* is denoted as Inertia(*M*) = {*n*_+_, *n*_−_, *n*_0_} where *n*_+_, *n*_−_, and *n*_0_ are the number of positive, negative, and zero eigenvalues of matrix M, respectively. One can prove that the reduced Hessian matrix is positive definite if Inertia(*M* ^𝓁^(*κ*_*w*_)) = {*n, m,* 0}, where we recall that *n* is the dimension of the variable vector *w* and *m* is the number of constraints. Notably, one can obtain the inertia of *M* ^𝓁^(*κ*_*w*_) without computing the eigenvalues of the matrix. This is done by using modern sparse symmetric factorization routines such as `MA57` or `Pardiso` [48]. Such routines factorize the matrix *M* ^𝓁^(*κ*_*w*_) as *LBL*^*T*^ where *L* is a lower triangular matrix and *B* is a matrix with 1 × 1 and 2 × 2 blocks in the diagonal. One can show that the number of positive and negative eigenvalues of *M* ^𝓁^(*κ*_*w*_) are the number of positive and negative eigenvalues of *B* (which are easy to determine).

Modern interior-point solvers are equipped with highly sophisticated safeguarding techniques that enable the solution of highly nonlinear problems. A powerful approach is called a filter line-search method, in which one seeks to find a step-size *κ* such that the trial Newton iteration *w*^𝓁 +1^ = *w*^𝓁^ + *κ*Δ*w*^𝓁^ either decreases the objective function or the constraint violation ||Π(*w*^𝓁^)||. If the step is accepted, the current values for the objective and constraint violation (Φ(*w*^𝓁^), Π(*w*^𝓁^)) are stored in a filter (a history of previous successful iterations). At the next iterate, one requires that the Newton step is not in the filter and that it improves either the objective or the constraint violation. This rather simple strategy is extremely effective in practice.

We highlight the fact that the proposed discretization approach bypasses the need to repetitively simulate the dynamic model (the discretized dynamic model contained in Φ(*w*) is solved progressively by Newton’s method). This brings substantial computational savings. Moreover, since the discrete-time model is solved at the solution *w*^*\**^ of the NLP (9), we have that the discrete-time states 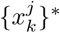 approximate the state trajectories *x*(*t*), *t ∈ 𝒯*_*k*_, *k ∈ 𝒦*. In the absence of inequality constraints, one can also show that the reduced Hessian *Z*^*T*^ *H*(*w*^*\**^, *λ*^*\**^)*Z* approximates the Hessian of the negative log-likelihood function *∇*_*θθ*_*L*(*θ*^*\**^) in a neighborhood of *w*^*\**^ (which contains *θ*^*\**^). We thus have that the reduced Hessian approximates the inverse parameter covariance matrix 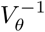. When inequality constraints are present, some of the parameters or state variables might hit their physical bounds and this deteriorates the approximation. When the parameters are not unique (the reduced Hessian has zero eigenvalues), the parameter covariance matrix is singular.

#### Structured Linear Algebra

A key advantage of using interior-point solvers is that they enable *modular linear algebra* implementations. For instance, the multi-experiment structure of problem (10) permeates down to the linear algebra system, to give a system of the form:

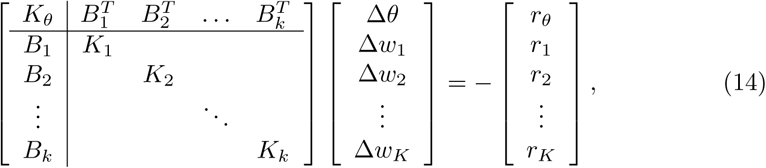

where Δ*θ* is the Newton step for the parameters and Δ*w*_*k*_ = (Δ*x*_*k*_, Δ*λ*_*k*_) is the Newton step for variables in experiment *k*. The above system is said to have a block-bordered diagonal (BBD) structure. Here, we have that:

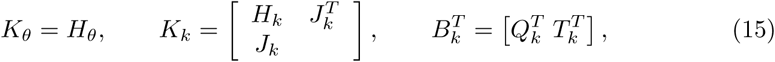

where 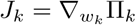, *T*_*k*_ = *∇*_*θ*_Π_*k*_, *H*_*θ*_ = *∇*_*θθ*_*ℒ* + *κ*_*w*_𝕀, *H*_*k*_ = *∇*_*w w*_ *ℒ* + *W*^−1^*V*_*k*_ + *κ*_*w*_𝕀, 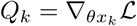, *r*_*θ*_ = *∇*_*θ*_*ℒ*, and 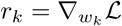.

The BBD matrix is a permutation of the augmented matrix *M* ^𝓁^(*κ*_*w*_) (obtained by ordering variables by experiment). The BBD matrix can thus be expressed as *P* ^*T*^ *M* ^𝓁^(*κ*_*w*_)*P* where *P* is a permutation matrix. The permutation does not affect the eigenvalues of the matrix. The BBD system (14) can be solved in parallel by using a Schur complement decomposition approach [45,50]. This requires the solution of the linear algebra systems:

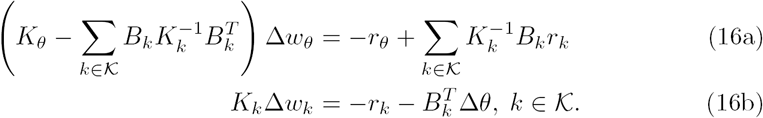

Here, 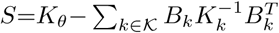 is the *Schur complement matrix* which has the same dimension as the number of degrees of freedom (in our case the number of parameters). The key observation is that the experiment matrices *K*_*k*_ can be factorized by using an *LBL*^*T*^ factorization (by using `MA57` or `PARDISO`) *in parallel*. As a result, Schur decomposition can achieve high computational efficiency in estimation problems with many experimental data sets.

When using a Schur decomposition, one can estimate the inertia of the BBD matrix by using Haynsworth’s formula:

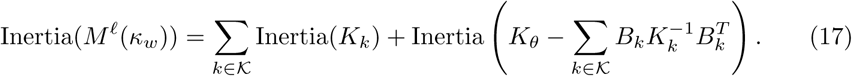

We recall that *n* = *n*_*θ*_ + ∑_*k*_ *n*_*k*_ and *m* = ∑_*k*_ *m*_*k*_. Consequently, if we have that Inertia(*K*_*k*_)={*n*_*k*_, *m*_*k*_, 0} for all *k ∈ 𝒦* then Inertia(*M* ^𝓁^(*κ*_*w*_)) = {*n, m,* 0} if and only if Inertia(*S*) = {*n*_*θ*_, 0, 0} (i.e., the Schur complement is positive definite). One can obtain the inertia of the blocks *K*_*k*_ and *S* using *LBL*^*T*^ factorization. This allows us to test observability of the parameters.

### Uncertainty Quantification

The estimation problem under the MAP framework gives the values of the parameters *θ*^*\**^ that maximize the parameter posterior density. However, a characterization of the entire posterior is necessary to assess parameter uncertainty. The posterior covariance may be approximated from the reduced Hessian at the solution of the problem *w*^*\**^ and the covariance matrix can be used to determine ellipsoidal level sets of the posterior (confidence regions). This approach, however, might fail to capture nonlinear and constraint effects [23]. In this work, we circumvent these issues by using a randomized maximum a posteriori (rMAP) approach. Under this method, the posterior distribution is explored by using random perturbations on the experimental data (which can be easily parallelized). The rMAP framework can also deliver *approximate samples* from the parameter posterior distribution and implicitly captures nonlinear and constraint effects. To show this, we use the implicit mapping representation 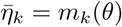. Under this representation, the posterior density (2) can be expressed as:

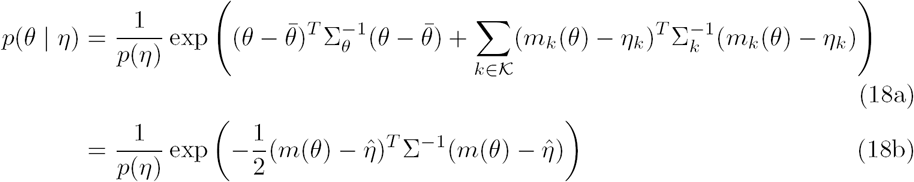

where *m*(*θ*) := (*θ, m*_1_(*θ*), *…, m*_*K*_(*θ*)), ∑ := diag(∑_*θ*_, ∑_1_, ∑_2_, *…,* ∑_*K*_), and 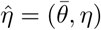. Here, we redefine 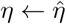 to enable compact notation. Since *θ*^*\**^ is a solution of the MAP problem, we have that:

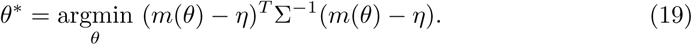

If the mapping *m*_*k*_(·) is continuously differentiable, we have that:

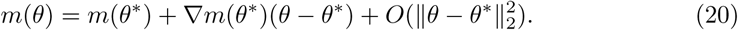

To enable compact notation we define *η*^*\**^ = *m*(*θ*^*\**^) and *∇m*^*\**^ = *∇m*(*θ*^*\**^). We have that *θ*^*\**^ satisfies the stationary condition of (19):

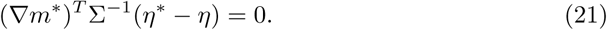

We use (20) to obtain a second-order Taylor approximation of the posterior as:

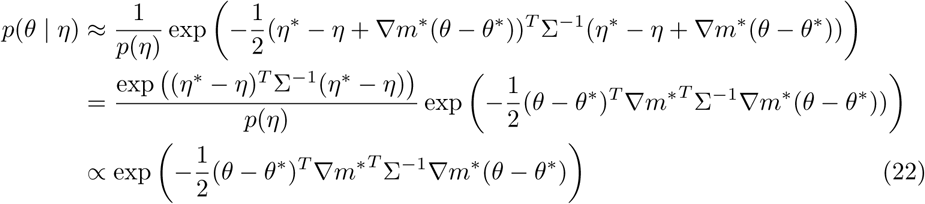

This implies that the posterior is *approximately* represented as:

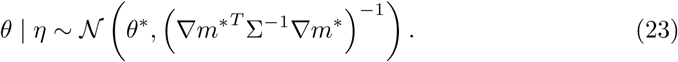

We recall that the output error is Gaussian and we can thus write *η* = *m*(*θ*) + *∈* with *∈ ∼ 𝒩* (0, ∑). We now consider the MAP problem with randomly perturbed data:

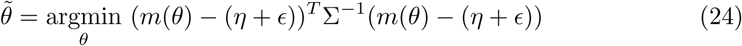

and note that

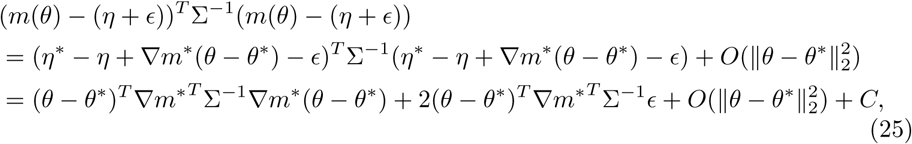

where *C* is a constant. Consequently, for sufficiently small *∈*, we can linearize the mapping *m*(·) to obtain an approximate solution of (24) of the form:

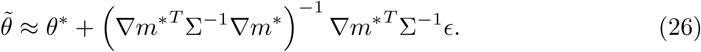

Here, we observe that the right-hand side of (26) is Gaussian with mean *θ*^*\**^ and covariance:

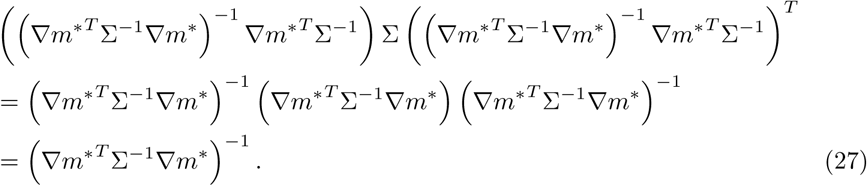

We thus have that:

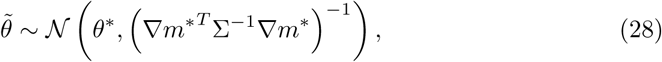

Consequently, solving (24) provides an approximate sample from the posterior distribution *p*(*θ| η*). The sampling procedure (24) is accurate up to second order. To obtain an exact sampling from the posterior, one needs to implement a rigorous MCMC scheme. The MCMC scheme removes the bias that appears in the rMAP sample density, which results from the second order approximation [34,36]. Several works in the literature, however, report that accurate posterior densities can be obtained using an rMAP scheme [35,51,52]. We also note that the rMAP scheme implicitly captures nonlinearities and physical constraints when computing samples from the posterior. In particular, solving the perturbed problem (24) corresponds to solving the MAP problem (1) with randomly perturbed data and the MAP problem enforces constraints and handles the full nonlinear model.

## Algebraic Modeling Platforms

Having an algebraic representation of the estimation problem has many practical and computational advantages. In particular, one can implement the estimation problem in easy-to-use and open-source modeling languages such as `JuMP` [53], `Plasmo.jl` [54], and `Pyomo` [55,56]. These modeling languages are equipped with automatic differentiation techniques that compute exact first and second derivatives. Derivative information is communicated to optimization solvers without any user intervention. Modern algebraic modeling languages such as `Plasmo.jl` and `Pyomo` also allow users to convey structural information to the solvers. This is beneficial in the case of parameter estimation, where the structure can be exploited to enable parallelism and the use of high-performance computing clusters. In our framework, we use the modeling language `Plasmo.jl` to express multi-experiment estimation problems as graphs. Our implementation using `Plasmo.jl` is illustrated in Fig 2. The full `Julia` script is available at https://github.com/zavalab/JuliaBox/tree/master/MicrobialPLOS. We highlight that the same script can be used to solve the estimation problem using a general NLP solver such as `Ipopt` on a single-processor computer or with a structure-exploiting parallel NLP solver such as `PIPS-NLP` on multiple parallel computing processors (this might be a multi-core computing server or a large-scale computing cluster). This allows users with limited knowledge on scientific computing to gain access to advanced high-performance computing capabilities.

**Fig 2.**
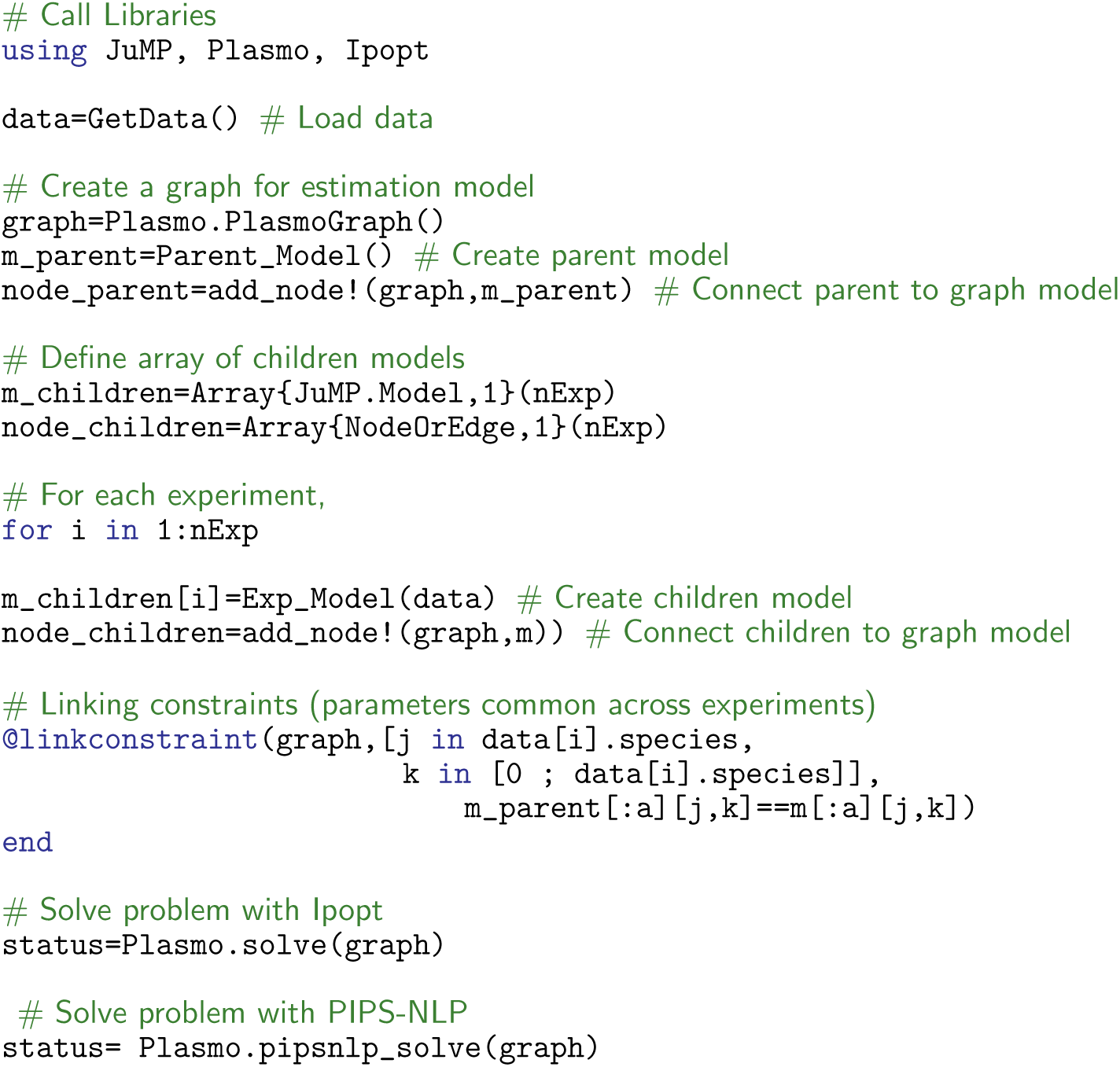
Snippet of parameter estimation implementation in Plasmo.jl.

## Results

Human gut microbial communities and microbiomes are highly dynamic networks coupled by positive or negative interactions and numerous feedback loops that display complex behaviors [57-60]. The generalized Lotka-Volterra (gLV) model provides a useful approach to capture such behavior [61-65]. Specifically, gLV captures single species growth rates and intra-species and inter-species positive and negative interactions. We apply the proposed NLP framework to estimate the growth and interaction parameters of the gLV model from experimental data collected in [60]. The microbial species involved in the experiments are shown in Table 1. Experiments were designed to study the synthetic ecology encompassing 12 prevalent human-associated intestinal species.

**Table 1.** List of microbial species used in case study.

| Label | Full name | Abbreviation |
| --- | --- | --- |
| 1 | *Blautia hydrogenotrophica* | BH |
| 2 | *Collinsella aerofaciens* | CA |
| 3 | *Bacteroides uniformis* | BU |
| 4 | *Prevotella copri* | PC |
| 5 | *Bacteroides ovatus* | BO |
| 6 | *Bacteroides vulgatus* | BV |
| 7 | *Bacteroides thetaiotaomicron* | BT |
| 8 | *Eggerthella lenta* | EL |
| 9 | *Faecalibacterium prausnitzii* | FP |
| 10 | *Clostridium hiranonis* | CH |
| 11 | *Desulfovibrio piger* | DP |
| 12 | *Eubacterium rectale* | ER |

The gLV model is given by:

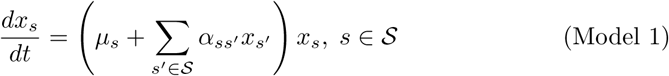

where *𝒮* = {1, 2, *…,𝒮*} is the set of microbial species, *x*_*s*_: ℝ → ℝ is the trajectory of the abundance of species *s∈ 𝒮, µ*_*s*_ is the growth rate of species *s*, and *α*_*ss’*_ is the interaction parameter that captures the effect of the abundance of species *s*^*’*^ on the growth rate of species *s*. Species *s* and species *s*^*’*^ are referred to as recipient species and donor species, resepectively.

The parameters (growth rates and interaction) cannot be calculated directly from first-principles and must be estimated from experimental data. The means of the prior densities of the parameters are assumed to be 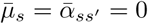 and their standard deviations are assumed to be 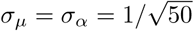. Such values are empirically determined by selecting the standard deviation values that give biologically feasible parameter estimates (the range of biologically feasible parameter values are 0.09 *< µ*_*s*_ *<* 2.1, −10 *< α*_*ij*_ *<* 10, and −10 *< α*_*ii*_ *<* 0). The variances for the output measurements are assumed to be *σ*_*k,s*_(*t*) = 0.05 max(0.1, *η*_*k,s*_(*t*)). There are a total of 156 parameters including 12 monospecies growth rate and 144 interaction parameters (12 x 12). The set of experiments 𝒦 includes 12 monospecies experiments and 66 pairwise community experiments (total of *K* = 78 experiments). The estimation problem contains a total of 144 differential equations (i.e. the model is a system of ordinary differential equations on ℝ^144^). The computational characteristics of the estimation problem are summarized in Table 2 (labeled as P1).

**Table 2.** Characteristic of estimation problems used in scalability studies.

|  | Original | Original+Synthetic | Synthetic |  |  |  |
| --- | --- | --- | --- | --- | --- | --- |
| Label | P1 | P2 | S1 | S2 | S3 | S4 |
| Number of Species | 12 | 12 | 12 | 24 | 36 | 48 |
| Number of Diff. Equations | 144 | 1,584 | 144 | 576 | 1,296 | 2,304 |
| Number of Parameters | 156 | 156 | 156 | 600 | 1,332 | 2,352 |
| Number of Experiments | 78 | 858 | 78 | 300 | 666 | 1,176 |
| Number of Data Points | 1,704 | 18,744 | 1,632 | 5,568 | 11,808 | 20,352 |
| Number of NLP Variables | 91,644 | 1,006,524 | 83,004 | 340,824 | 773,460 | 1,380,912 |
| Number of NLP Constraints | 91,488 | 1,006,368 | 82,848 | 340,224 | 772,128 | 1,378,560 |

Problem P1 includes the original experimental data [60] for a 12-species microbial community (consisting of 12 mono-species and 66 pairwise community experiments). The sampling frequency and the experiment duration of the mono-species experiments are 30 minutes and 24 hours, respectively. The sampling frequency in the pairwise community experiments is 12 hours and the experiment duration range from 60 to 72 hours. In the pairwise experiments, the media are diluted by 1/20 once every 24 hours. The dynamic model is discretized using an implicit Euler scheme with 5 equally-spaced discretization points (monospecies experiments) and 120 equally-spaced discretization points (pairwise experiments). The data used in P2 includes the original data of P1 and 10 additional synthetic data sets obtained with random data perturbations. Data for problems S1-S4 is synthetic and is obtained by running a simulation of the community model with fixed parameters and by adding 5% noise to the outputs. The data sampling characteristics (e.g., frequency, duration, dilution patterns) are the same as those of P1. The parameter values used for simulations are randomly generated from *µ*_*s*_ *∼ 𝒩* (0.3, 0.1^2^), *α*_*ss*_ *∼ 𝒩* (−1, 0.1^2^), and *α*_*ss’*_ *∼ 𝒩* (0, 0.1^2^).

### Observability and Regularization

Parameter observability was checked by solving the MAP formulation for P1 (which uses the available experimental data) and by checking the inertia of the augmented system at the solution (reported by `Ipopt`). Here, we omitted the prior regularization term *ϕ*(·). We found that parameters obtained from P1 are *not unique* (not observable from the available data). Moreover, we found that the estimated parameter values without regularization have unrealistic (non-physical) values, see Fig 3 (a). This observation justifies the need to use prior information. The results obtained by adding L1 and L2 priors to the MAP formulation are presented in Fig 3 (b,c). Unique parameter estimates were found when L1 or L2 priors were used. We also found that the L1 prior induces sparser solutions (many parameters are zero). For the remainder of the results, we use the formulation with an L2 prior.

**Fig 3.**
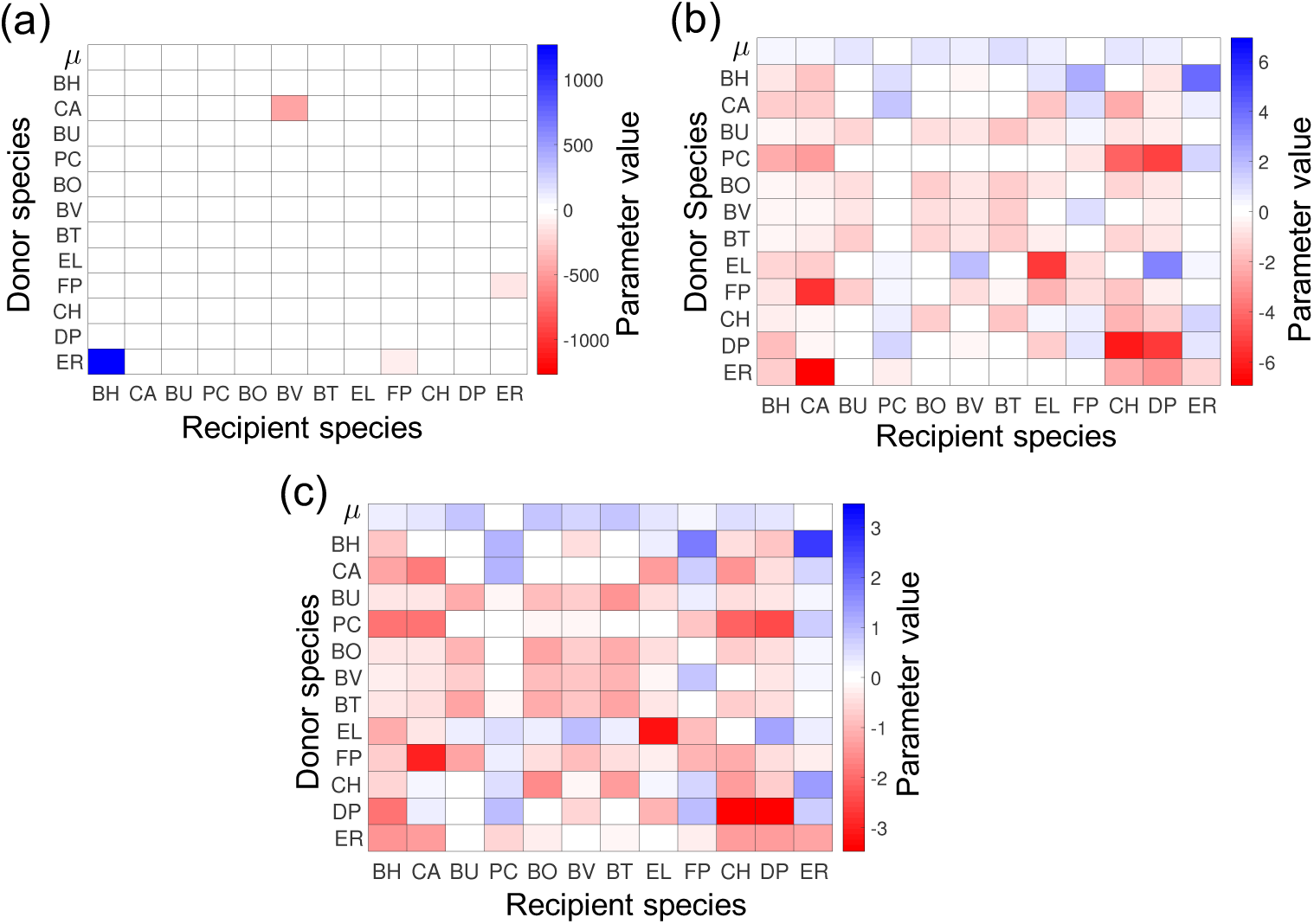
Parameter estimates with MAP formulations (Model 1). **(a)** Estimates using MAP formulation with no prior information. **(b)** Estimates using L1 prior. **(c)** Estimates using L2 prior. (a-c) The first row shows values for the growth rate parameters *µ*_*s*_ and the rest of the rows show values for the interaction parameters *α*_*ss’*_(·) The species name corresponding to *s* and *s*^*’*^ are presented on the *x* and *y* axes. Recipient and donor species are on the *x* and *y*-axis, respectively.

## Model Fitting

Model validation was performed by assessing the goodness of fit to the experimental data (Fig 4). We can see that the model is capable of fitting most of the data points, but there are a number of experiments where the model prediction deviates significantly from the experimental data (such experiments are highlighted with red boxes). Furthermore, we can observe outliers at single data points (highlighted with red circles). Poor fitting can be caused by either bad local minima (the optimization solver finds a local optimal solution rather than the global optimal solution) or by a structural model error (the model structure is incapable of capturing the actual behavior of the system). To avoid bad local minima, we solved the MAP problem with multiple starting points. Such an approach increases the probability to find the global optimum, but obtaining a rigorous certificate of a global minimum is computationally challenging (rigorous global optimization techniques are currently not scalable to large problems). We found that the use of multiple starting points does not improve the model fit. Consequently, we attribute fitting errors to the model structure itself. In particular, the gLV model neglects various physical and biological phenomena such as lag phase or interaction coefficients that change as a function of time [60]. To investigate structural errors, we solved the MAP problem with a variant of the gLV model. In particular, we investigated the saturable gLV model [66,67] (Model 2):

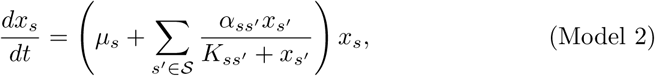

**Fig 4.**
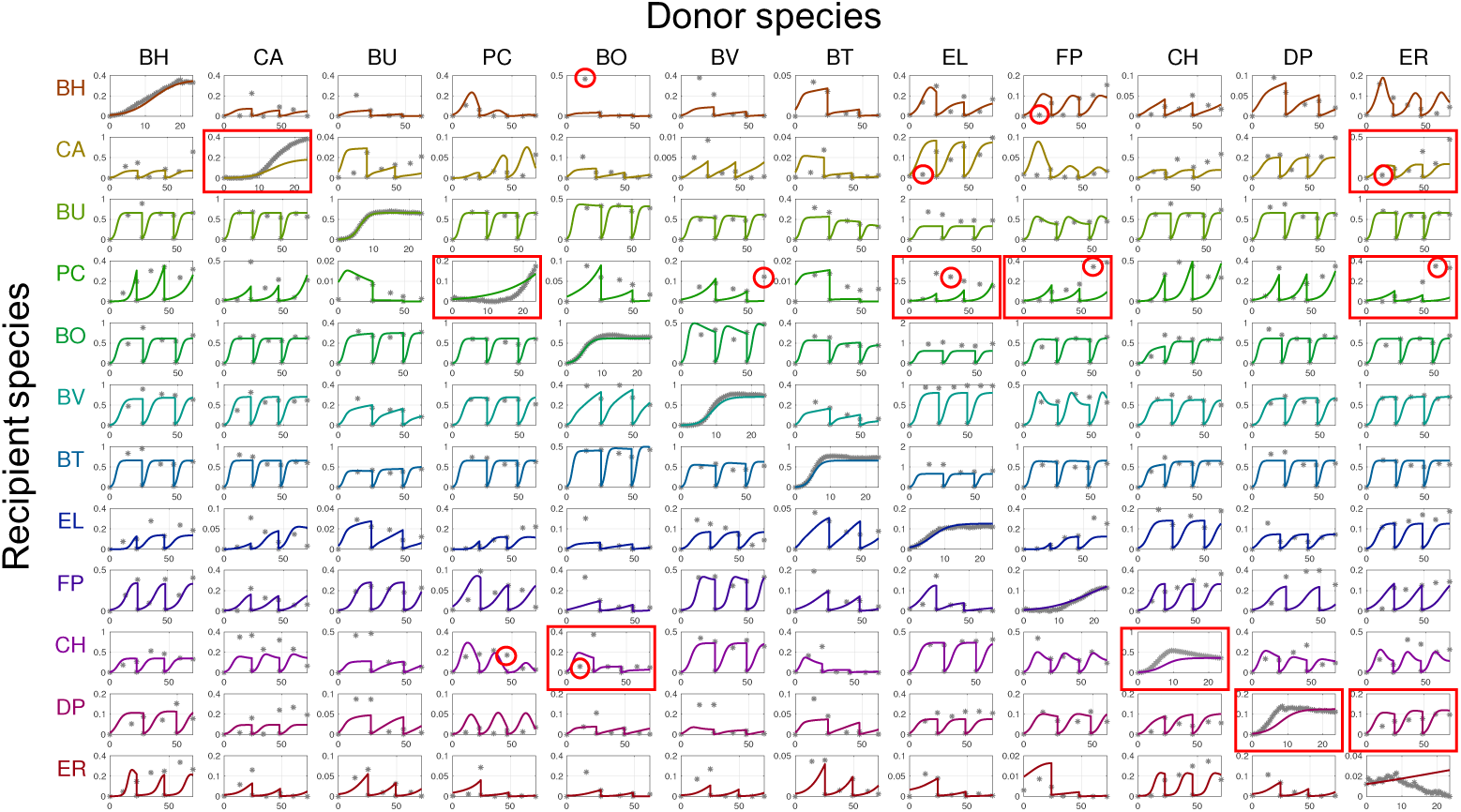
Fitting of experimental data (Model 1). Subplots show the measured and predicted species abundance in the microbial community. The subplots on the diagonal show fitting for mono-species experiments. Subplots on the *i*-th row and *j*-th column shows fitting for the corresponding pairwise culture (the abundance of species *i* in the presence of species *j*). Recipient and donor species are listed in rows and columns, respectively. For each subplot, the *y*-axis represents the absolute abundance in the community based on relative abundance multiplied by total biomass (OD600) and the *x*-axis represents the experiment time in hours. Data points are denoted by grey dots and dynamic model trajectories are denoted by solid lines. The data points highlighted with red circles are data points corresponding to the ten largest errors 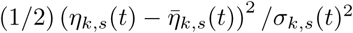. The subplots highlighted with red boxes are subplots for the experiments with the ten largest total prediction errors 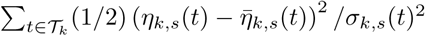

where *K*_*ss’*_ *>* 0 are additional interaction parameters. The saturable model exhibits a higher degree of nonlinearity than the gLV model and includes *S*^2^ more parameters (the number of degrees of freedom increases from *S*^2^ + *S* to 2*S*^2^ + *S*). As a result, the saturable gLV model provides more flexibility to improve model fitting. The model fitting obtained with the saturable gLV form is illustrated in Fig 5. As can be seen, significant improvements; in particular, the overall fitting error

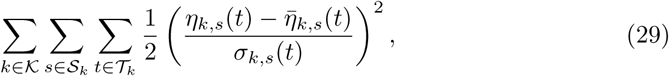

**Fig 5.**
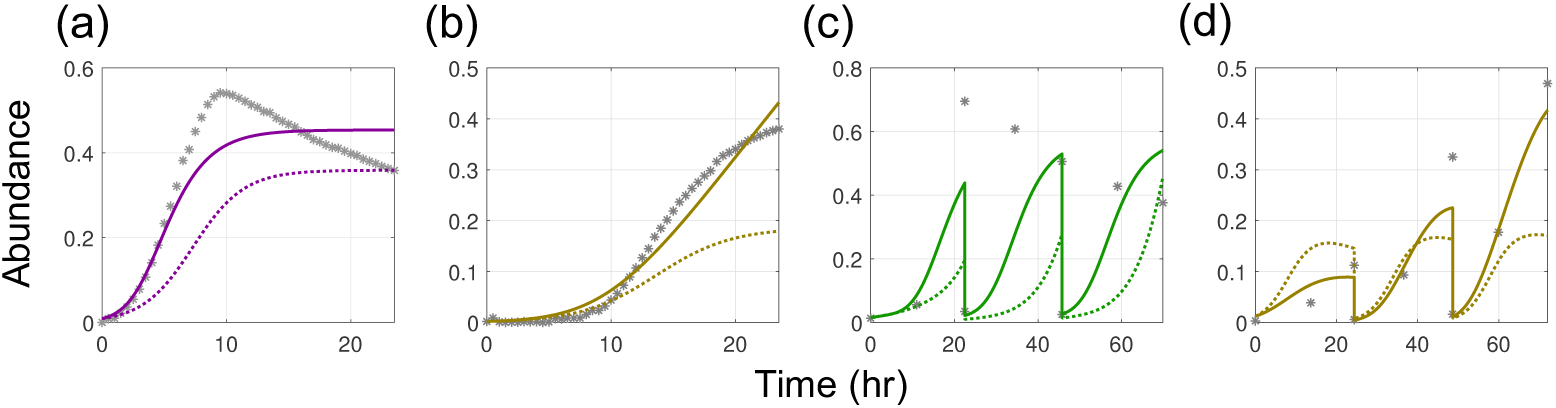
Improvement of model fitting with saturable gLV model (Model 2). Model fitting for 4 experiments selected among the experiments with 10 largest total prediction errors. The gLV model fits (dotted line) are compared with those of the saturable model (solid line). **(a)** Model fits to monospecies experiment with CH. **(b)** Model fits to monospecies experiment with CA. **(c)** Model fits to pairwise community experiment with PC in the presence of EL. **(d)** Model fits for pairwise community experiment with CA in the presence of ER.

was reduced by 30%. Increasing the number of degrees of freedom can cause overfitting, however, and this can make the model less predictive. Consequently, there is a trade-off between fit and predictability.

Bayesian information criteria (BIC) is an approach for model selection [68-70]. BIC can be used as a score that represents the likelihood of the model that considers not only model fitting but also the number of parameters. In particular, the model with the smallest BIC can be considered as the most likely model. In BIC, one aims to compare the posterior probability of the models given the experimental observation. The posterior probabilities can be indirectly compared by comparing the marginal probability *p*(*η*) = ∫ *p*(*η θ*)*p*(*θ*)*dθ* of the experimental observation for each model. By applying a Laplace approximation, one can derive a reasonable approximation of −2 log *p*(*η*) as follows.

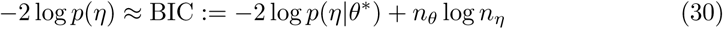

Recall that *n*_*η*_ is the dimension of the output vector *η* (the number of experimental observations) and *θ*^*\**^ is the optimal parameter estimate. The log-likelihood term –2 log *p*(*η| θ*^*\**^) corresponds to the prediction error (the squared errors across organisms and data sets) value of the MAP problem and is obtained while solving the problem. As can be seen from (30), the number of parameters are penalized in the score. Thus, the effect of overfitting that comes from increasing the number of parameters can be prevented. Our results indicate that the BIC of the gLV model is 44, 798 and the BIC of the saturable model is 31, 572. This implies that the saturable model is the more likely model based on the available experimental data. We highlight, however, that the Laplace approximation assumes independent and identically distributed sampling and a sufficient number of samples (which does not strictly hold for our case). Model selection based on BIC with specialized experimental data is an interesting direction of future work.

To mitigate outliers, we solved the MAP problem with a CVaR norm and *β* = 0.9 (to penalize the 10% largest errors). The model fitting obtained with the CVaR formulation is shown in the supporting information and relevant results are summarized in Fig 6. It can be observed that the fitting errors for the outliers obtained with the standard MAP formulation are reduced. The effect of the CVaR formulation is also evident when analyzing the prediction error histogram (see Fig 7). In particular, we observe that the tail of high prediction errors becomes less pronounced under the CVaR formulation. In particular, the mean of the 10% largest errors decreases by 18% (from 167.81 to 137.04). On the other hand, it can also be observed that the mean error increases under CVaR and that the tail of small errors shrinks. This illustrates the fundamental trade-off that usually arises in robust statistics. The behavior induced with CVaR aids estimator performance because it prevents overfitting experimental data sets.

**Fig 6.**
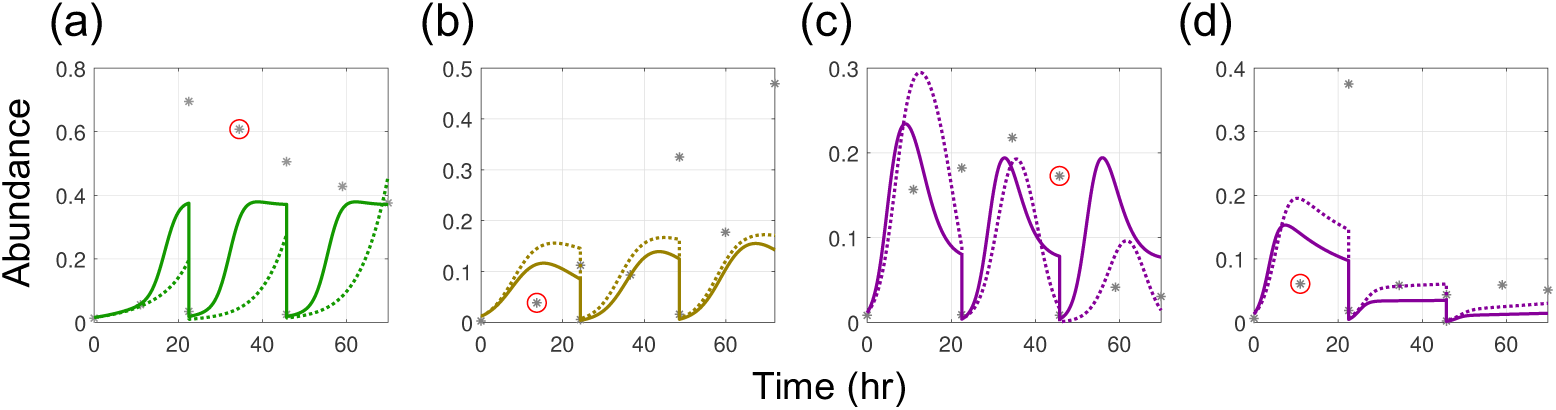
Improvement of model fitting by using CVaR (*k*-max) MAP formulation (Model 1). Model fits for 4 experiments selected among the experiments with 10 largest prediction errors 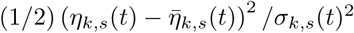. The model fits from standard MAP formulation (dotted line) are compared with the model fits from CVaR formulation with *β* = 0.9 (solid line). **(a)** gLV model fits to pairwise community experiment of PC in the presence of EL. **(b)** gLV model fits to pairwise community experiment of CA in the presence of ER. **(c)** gLV model fits to pairwise community experiment with CH in the presence of PC. **(d)** gLV model fits to pairwise community experiment with CH in the presence of BO.

**Fig 7.**
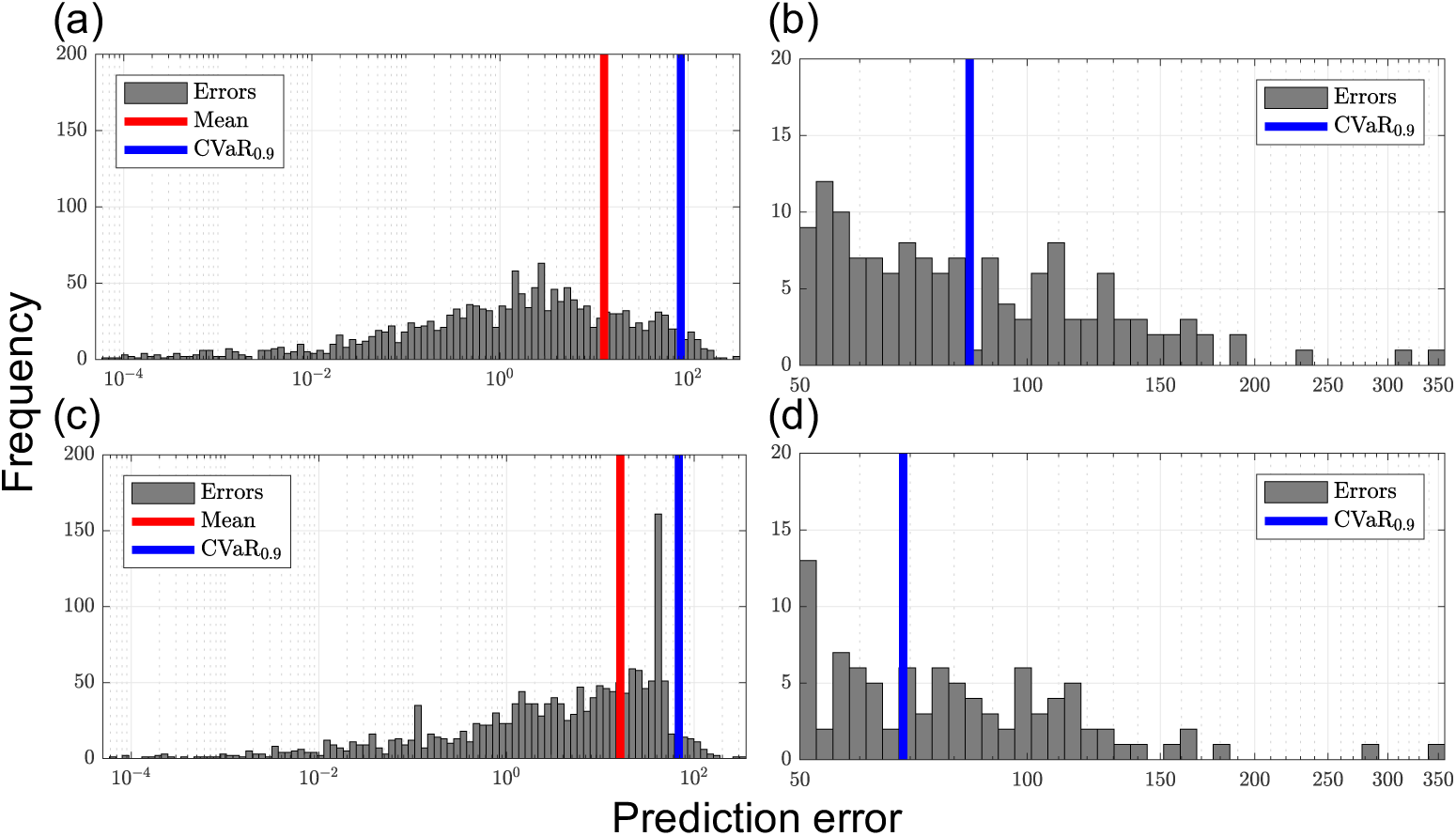
Histograms of fitting errors (Model 1). **(a)** Error histogram for the standard MAP formulation. **(b)** Tail region of (a). **(c)** Error histogram for CVaR formulation. **(d)** Tail region of (c). (a-d) The *x*-axis represents the value of prediction error evaluated at the solution and the *y*-axis represents the frequency. The red and blue line represent quantiles: the overall mean of prediction errors (red) and the mean of largest 10% errors (blue).

### Inference (Posterior) Analysis

We used rMAP to assess the uncertainty of the 156 parameters estimated from P1 using the available experimental data. To do so, we draw data perturbations as *η*_*k,s*_(*t*) ← *η*_*k,s*_(*t*) + *∊*_*k,s*_(*t*) with *∊*_*k,s*_(*t*) ∼ 𝒩 (0, *σ*_*k,s*_(*t*)^2^). We solved 500 MAP problems to obtain parameter samples and use this to approximate the covariance matrix for the posterior. The marginals for the posterior are shown in Fig 10 and the standard deviations are shown in Fig 8 (a). A large standard deviation indicates that the estimated parameter value is not reliable. We note that about half of the parameters can be estimated reliably while the other half exhibit significant uncertainty. This indicates that more experimental data should be obtained. From the sample covariance, we generated 95% ellipsoidal confidence regions for each pair of parameters. The correlation plots of *µ*_*s*_ against *α*_*ss’*_ for *s, s*^*’*^ ∈ *𝒮* are shown in Fig 11 and the Pearson correlation coefficients are shown in Fig 8 (b). In an ideal case, the parameters should be uncorrelated because data should be sufficient to estimate each parameter reliably. Using our data set, however, we can observe strong correlations between the parameters *µ*_*s*_ and *α*_*ss*_ in Fig 11, and strong positive and negative correlations can also be found in Fig 8 (b).

**Fig 8.**
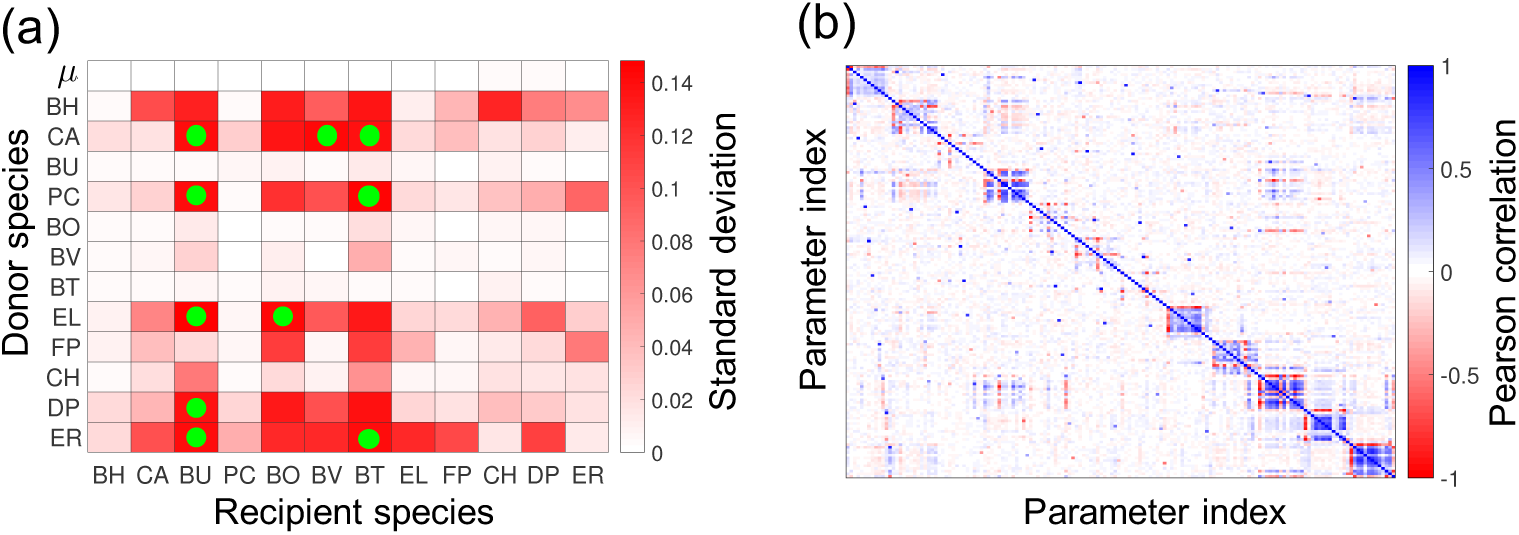
Parameter uncertainty and correlations (Model 1). **(a)** Heat map presents the standard deviation for the parameter posterior density. The first row shows the standard deviations of the growth rate parameters *µ*_*s*_ and the rest of the rows show the standard deviations of the interaction coefficients *α*_*ss’*_. Recipient and donor species are on the *x* and *y*-axis, respectively. The data points highlighted with green circles are data points corresponding to the ten largest standard deviations. **(b)** The heat map represents the Pearson correlation coefficients of the poterior distributions. The *x*-axis and the *y*-axis represents the index of parameters where the parameter vector is constructed as *θ* = (*µ*_1_, *α*_11_… *α*_1*S*_,*…, µ*_*S*_, *α*_*S*1_,*… α*_*SS*_). The block on the *i*-th row and *j*-th column is the Pearson correlation between the *i*-th and *j*-th component of parameter vector *θ*.

**Fig 9.**
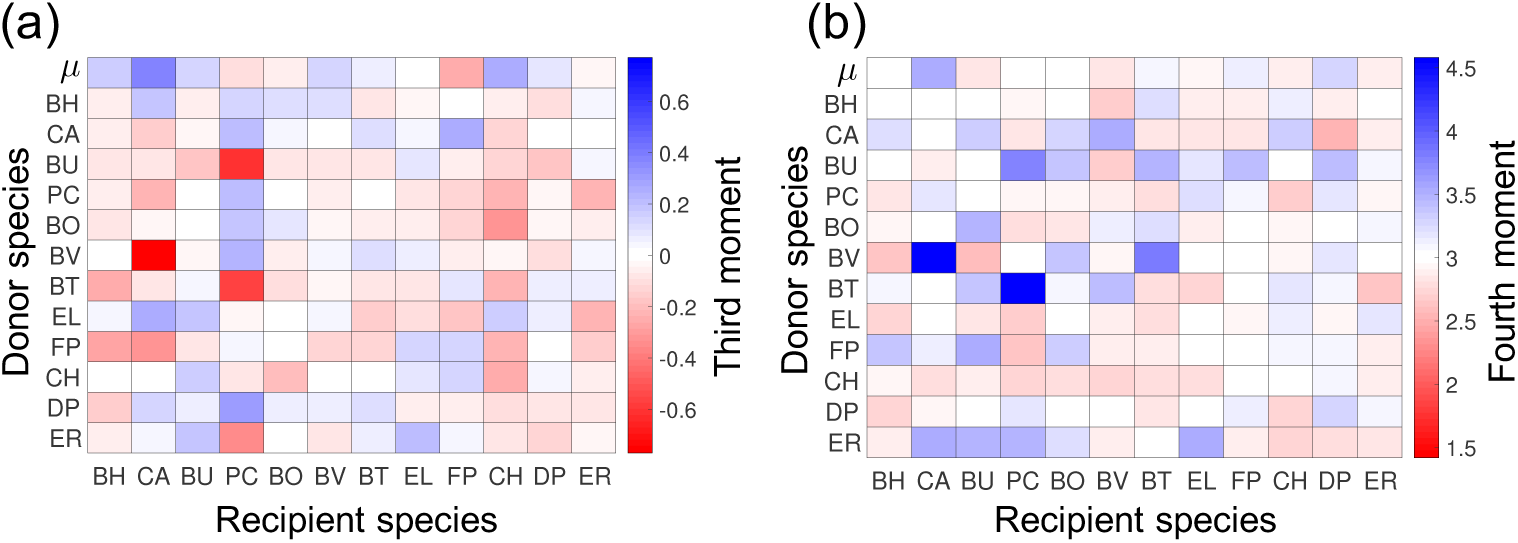
Third and fourth momentum of posterior (Model 1). **(a)** Heat map presents the third momentum of the parameter posterior density (normalized by *σ*^3^).**(b)** The heat map represents the fourth momentum of the poterior density (normalized by *σ*^4^). (a-b) The first row shows the standard deviations of the growth rate parameters *µ*_*s*_ and the rest of the rows show the standard deviations of the interaction coefficients *α*_*ss’*_. Recipient and donor species are on the *x* and *y*-axis, respectively.

**Fig 10.**
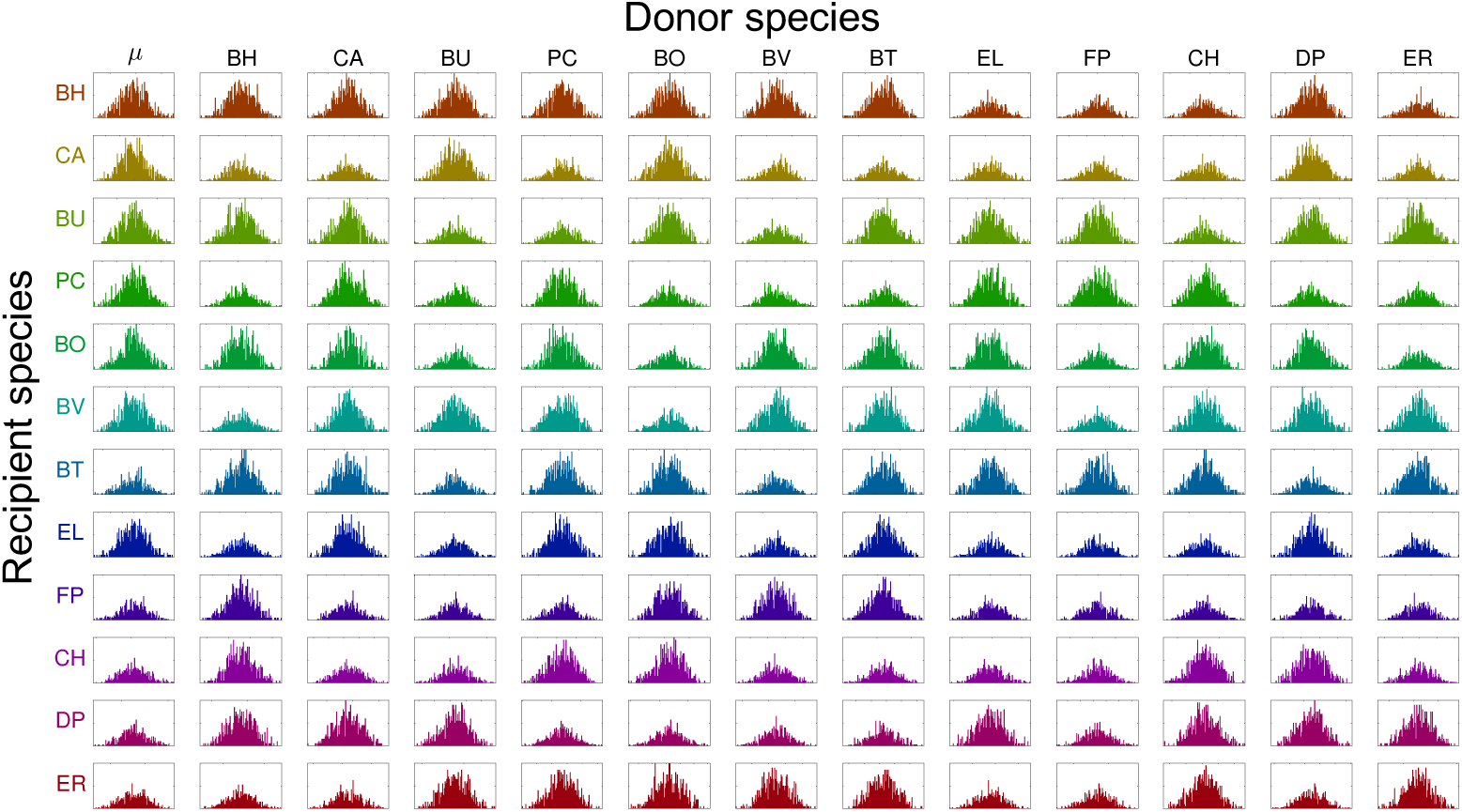
Posterior (marginal) densities for estimated parameters (Model 1). Each subplot shows the histogram of the samples from the approximate parameter posterior. The *x*-axis represents the values of the estimated parameters and the *y*-axis represents the frequencies. The subplots on the first column show the distribution of the growth rates *µ*_*s*_ and the rest of the subplots show distributions of the interaction parameters *α*_*ss’*_. Recipient and donor species are listed in rows and columns, respectively. The *x*-axis is scaled to show *µ ±* 3*σ* where the *µ* is the mean and the *σ* is the standard deviation of the posterior distribution.

**Fig 11.**
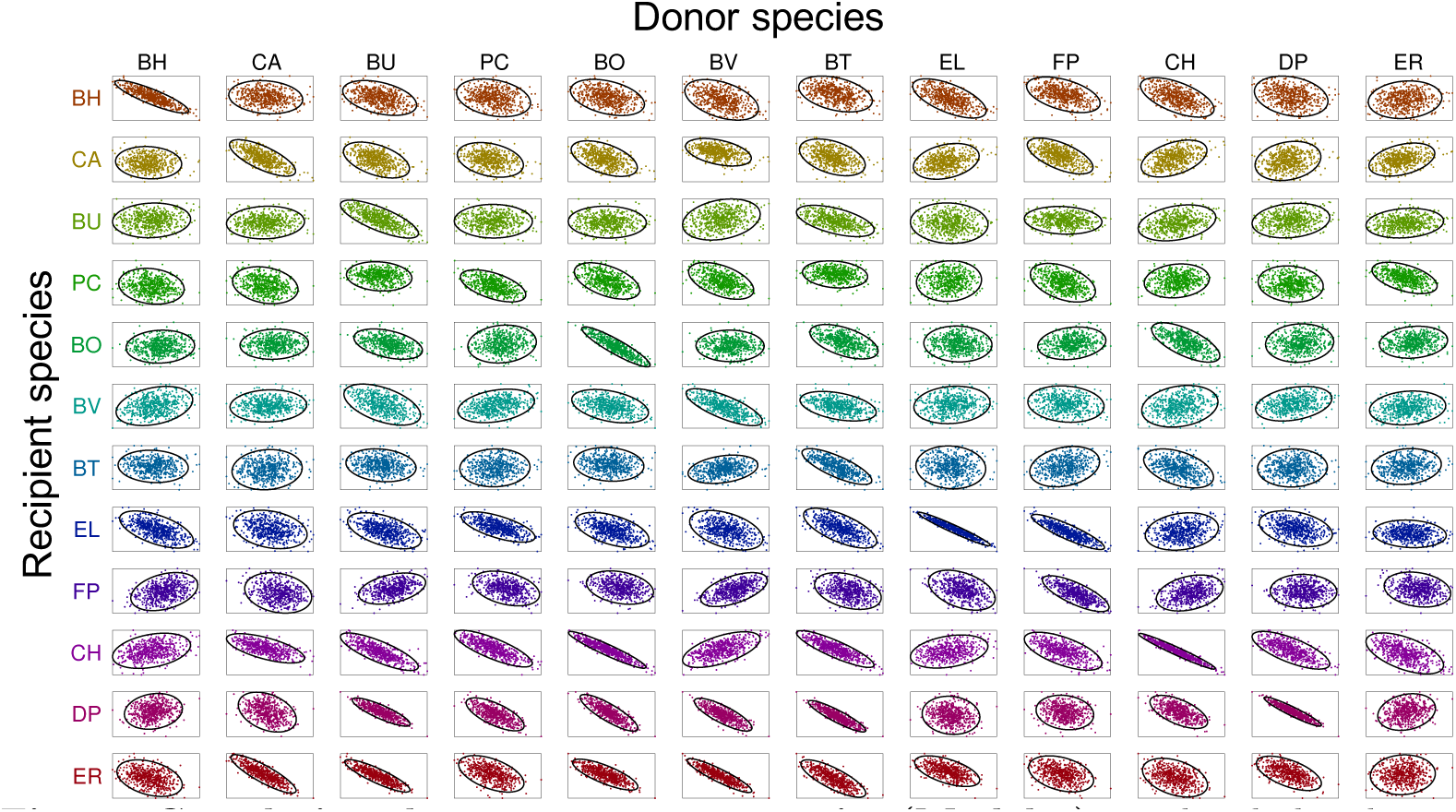
Correlations between parameter pairs (Model 1). Each subplot shows the 95% confidence regions (solid ellipses) of the approximate parameter posterior distributions and the sample points (dots). The subplots on the *s*-th row and *s*^*’*^-th column show the correlation of *µ*_*s*_ and *α*_*ss’*_. Recipient and donor species are listed in rows and columns, respectively. Only a representative subset of parameter pairs is presented (there are a total 12,090 pairs).

Furthermore, since the whole approximate distribution is obtained in the inference analysis based on rMAP framework, we can perform more sophisticated analysis on the characteristics of the distribution. In particular, one can investigate third and fourth moments (Fig 9) to examine the skewness and the kurtosis of the distribution. Such information can be used to investigate the deviation of the posterior distribution from the normal distribution. If the posteriors are stritly normally distributed, the third and fourth moments should be zero and 3*σ*^4^, respectively. However, we can observe that many posterior distributions deviate from such expectations. Thus, we can see that some of the distributions are not close to the normal distribution.

### Computational Scalability

We assessed the computational scalability of the estimation framework by analyzing problems with different sizes and characteristics. Problem P1 was implemented in the algebraic modeling platform `JuMP` and solved with the NLP solver `Ipopt` configured with the sparse linear solver `MA57`. Problem P1 with gLV model and L2 prior was solved in 134 seconds and 78 NLP iterations on a standard computing server with an Intel(R) Xeon(R) CPU E5-2698 v3 processor running at 2.30GHz. Problem P1 with gLV model and L1 prior was solved in 219 seconds and 68 NLP iterations with the same hardware. A comparable problem requires over 7 hours to solve using a simulation-based approach implemented in `Matlab` and that uses finite differences to obtain first derivatives [60]. Despite the significant gains in computational performance obtained with `Ipopt`, its solution time scales nearly quadratically with the number of data sets. To overcome this scalability issue, we compared the performance of the serial solver `Ipopt` against that of the parallel solver `PIPS-NLP` (which uses a Schur complement decomposition to perform linear algebra operations). To test the scalability of `PIPS-NLP`, we generated a larger version of the estimation problem (labeled as P2). This problem is created by adding synthetic data sets. The NLP corresponding to P2 has over one million variables and constraints (but the number of parameters is the same as that of P1). This problem was implemented in `Plasmo.jl`. The benefit of using a parallel approach is clearly seen in Fig 12. Here, we highlight that `PIPS-NLP` solved P2 in less than 10 minutes and 94 NLP iterations using 16 cores while `Ipopt` requires around 30 minutes and 67 iterations. Furthermore, `IPOPT` found a different local solution and the solution from `PIPS-NLP` had a better objective value. Fig 12(b) also shows that `PIPS-NLP` achieves nearly perfect strong scaling (speedup increases linearly with the number of cores).

**Fig 12.**
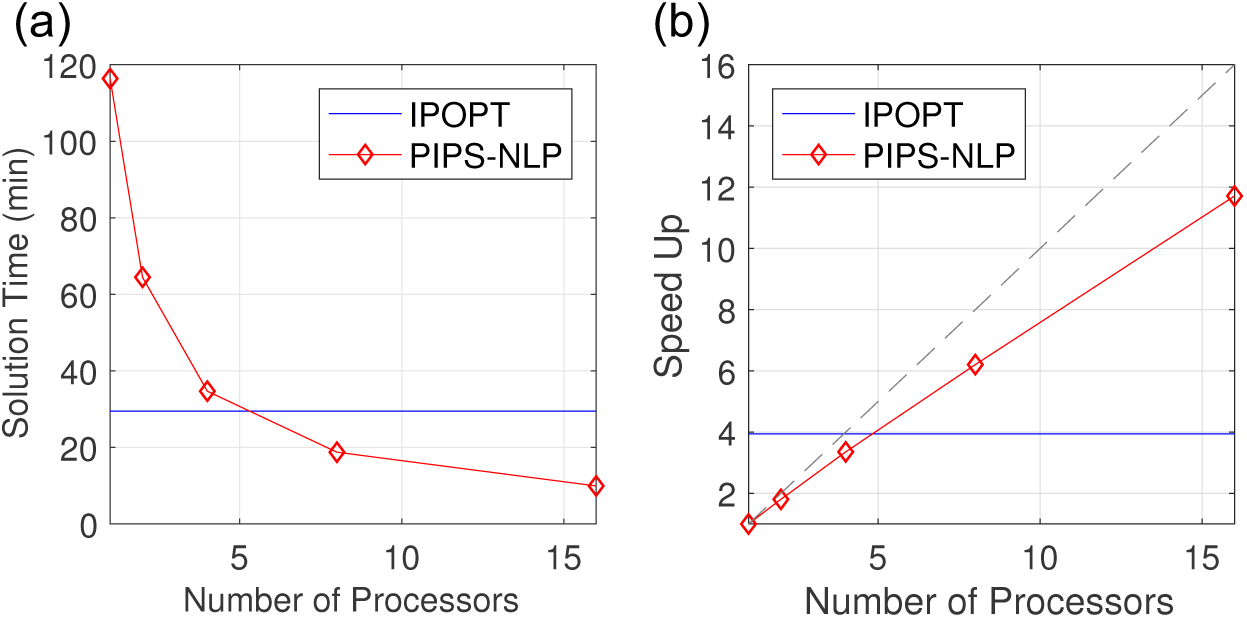
Performance comparison of general solver Ipopt and the structure-exploiting solver PIPS-NLP (Model 1). **(a)** Solution time for P2 using `Ipopt` and `PIPS-NLP`. The *y*-axis shows the solution time and the *x*-axis shows the number of cores used. For `Ipopt` the single core solution time is given by the horizontal blue line. **(b)** The *y*-axis represents the speed-up (the single-core solution time divided by the multi-core solution time). The blue line is the single-core solution time of `PIPS-NLP` divided by the single-core solution time of `Ipopt`. The grey dashed line represents the strong scaling line.

In rMAP-based uncertainty quantification, the main computational challenge was the repetitive solution of the optimization problems. However, such challenge can be overcome by using the existing solution information. The required number of iterations in NLP solver can be greatly reduced when a good starting point (initial guess of the solution) is available (often referred to as *warm start*). Since only small modifications are made to the original problem to formulate the rMAP problem, the NLP solution of rMAP problem is very similar to that of the original problem. Thus, by warm-starting the NLP with the original NLP solution, the computational efforts to solve rMAP problem can be significantly reduced. In particular, most rMAP sampling problem was solved in less than 10 NLP iterations while the original problem required NLP 78 iterations.

We also assessed computational capability in estimation problems with the larger number of species in the microbial community (which increases the number of differential equations and parameters). Here, we generated synthetic data using simulations for larger communities. The generated data are summarized in Table 2. The number of the parameters and of data points scales nearly quadratically with respect to the size of the community. The computation times are shown in Fig 13. The results indicate that, by using `PIPS-NLP`, one can solve estimation problems with up to 48 species in *less than 15 minutes and 40 NLP iterations* (using 12 parallel computing cores). We highlight that, to the best of our knowledge, problem S4 is the largest estimation problem reported in computational biology literature. This problem contains 2,304 differential equations, 2,352 parameters, and 20,352 data points. The corresponding NLP contains 1.3 million variables and constraints.

**Fig 13.**
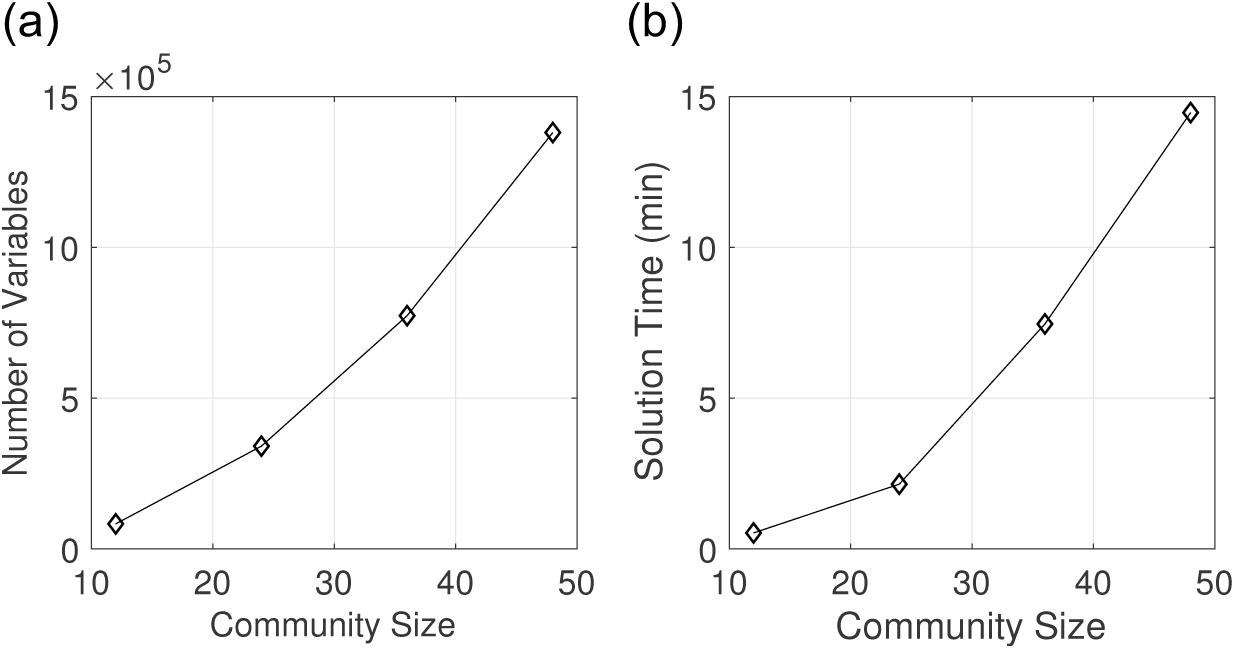
Computational scalability with larger communities (Model 1). **(a)** Number of variables against community size (total number of species). **(b)** The computation times for problems S1-S4 (see Table 2). The problems were solved with `PIPS-NLP` on 12 parallel cores (Intel(R) Xeon(R) CPU E5-2698 v3 processor running at 2.30GHz).

### Concluding Remarks

The high computational efficiency achieved with the proposed framework can enable kinetic modeling of complex biological systems ranging from biomolecular networks to high-dimensional microbial communities [71]. Indeed, the proposed framework can be used to construct and analyze high-fidelity models of whole-cells or microbiomes [72,73]. In particular, these methods can be applied to develop predictive dynamic models of multi-gene synthetic circuits interacting with host-cell processes for accurately predicting cell growth and synthetic circuit activity [74] or kinetic models of metabolite transformations driving community dynamics. These methods will advance our capability of integrating mechanistic modeling frameworks with large-scale experimental data. Furthermore, uncertainty quantification and observability analysis can provide valuable information to guide and accelerate experimental data collection. These capabilities are also essential in diagnosing structural model errors. The proposed framework uses state-of-the-art and easy-to-use modeling and solution tools that can be broadly applied to diverse biological systems and accessible to a wide range of users. Together, these advances will ultimately transform biology into a predictive and model-guided discipline.

